# *Sciara coprophila* larvae upregulate DNA repair pathways and downregulate developmental regulators in response to ionizing radiation

**DOI:** 10.1101/2021.10.28.466123

**Authors:** John M. Urban, Jack R. Bateman, Kodie R. Garza, Julia Borden, Jaison Jain, Alexia Brown, Bethany J. Thach, Jacob E. Bliss, Susan A. Gerbi

**Affiliations:** Brown University Division of Biology and Medicine, Department of Molecular Biology, Cell Biology and Biochemistry, Providence, RI 02912; Biology Department, Bowdoin College, Brunswick, ME 04011; Department of Embryology, Carnegie Institution for Science, Howard Hughes Medical Institute Research Laboratories, 3520 San Martin Drive, Baltimore, MD, 21218

## Abstract

Robust DNA damage prevention and repair strategies are crucial to faithful reproduction and inheritance of the genetic material. Although many molecular pathways that respond to DNA damage are well conserved through evolution, the quality and effectiveness of these systems can vary between species. Studies dating back for nearly a century document that the dark-winged fungus gnat *Sciara coprophila* (Order: Diptera; sub-order: Nematocera) is relatively resistant to irradiation-induced mutations that cause visible phenotypes when compared to the fruit fly *Drosophila melanogaster* (Order: Diptera; sub-order: Brachycera). However, the molecular responses to irradiation for *S. coprophila* have yet to be analyzed. To address this gap, we first characterized the effects of ionizing radiation on *S. coprophila* throughout its life cycle. Our data show that developing *S. coprophila* embryos are highly sensitive to even low doses of gamma-irradiation, whereas larvae can tolerate up to 80 Gy and still retain their ability to develop to adulthood with a developmental delay of 5 to 8 extra days in the larval stage. To survey the genes involved in the early transcriptional response to irradiation, we compared RNA-seq profiles of larvae with and without radiation treatment. Our analysis showed that 327 genes are differentially expressed in irradiated larvae, with 232 genes upregulated and 95 genes downregulated relative to controls. The upregulated genes were enriched for DNA damage response genes, including those involved in DNA repair, cell cycle arrest, and apoptosis, whereas the down-regulated genes were enriched for developmental regulators, consistent with the developmental delay observed in irradiated larvae. Thus, our study has laid the groundwork to further dissect how *Sciara* copes with radiation-induced damage.

## Introduction

Maintenance of genome integrity is central to an organism’s survival and its ability to faithfully pass genetic information to its offspring. Loss of genome stability and DNA mutation can result from errors in endogenous cellular events such as DNA replication and chromosome segregation, and from exposure to exogenous environmental agents that can alter or damage DNA. Living systems have therefore evolved various biochemical and developmental pathways that recognize and respond to errors and/or damage in their genetic code. In humans, altered function in these pathways is a common feature in the development and progression of cancer, allowing the accumulation of further mutations in a cancer lineage. A better understanding of diverse responses to DNA damage may therefore be helpful in developing novel approaches to disease.

High-energy ionizing radiation, including X- and gamma-rays, is an abundant and well-studied DNA damaging agent. Ionizing radiation can directly interact with DNA, causing lesions to individual bases, single-strand breaks (SSBs), and double-strand breaks (DSBs) (Han and Yu 2010). Alternatively, radiation can interact with other molecules in the cell to generate reactive oxygen species (ROS), which can themselves cause DNA damage including single base lesions and SSBs. In metazoans, DNA damage can be repaired by several well-characterized pathways, including base excision repair (BER), nucleotide excision repair (NER), mismatch repair (MMR), homology directed repair (HDR), and nonhomologous end-joining (NHEJ) (Chatterjee and Walker 2017). In addition, extensive DNA damage can trigger apoptotic pathways to remove cells with potentially unchecked genome instability.

While all living systems appear to have some ability to respond to damaged DNA, several species have evolved highly robust responses to ionizing radiation that stem from expanded DNA repair or DNA protection mechanisms. For example, the extremophile bacterium *Deinococcus radiodurans* can endure doses of gamma-rays several orders of magnitude higher than a typical mammal, in part due to their ability to reassemble highly fragmented genomic DNA into an intact chromosome via specialized synthesis and recombination pathways (Bentchikou *et al.* 2010). Similarly, tardigrades are highly resistant to extreme conditions such as desiccation and exposure to radiation, and several species encode a tardigrade-specific nuclear protein called Damage Suppressor (Dsup) that binds to nucleosomes and protects them from DNA damage induced by free radicals (Chavez *et al.* 2019). Notably, Dsup expression in culture human cells can increase their resistance to X-irradiation (Hashimoto *et al.* 2016), demonstrating that radiation-protective mechanisms that evolve in one species have the potential to function more broadly.

Historically, ionizing radiation has been an important tool for introducing mutations in model organisms for genetic studies, and was central to our understanding of the mutability of genes through Muller’s experiments in *Drosophila melanogaster* and to the development of the ‘one gene, one enzyme’ hypothesis in *Neurospora crassa* (Carlson 2013; Strauss 2016). Treatment of model organisms such as *Drosophila* and *Neurospora* with moderate doses of radiation creates diverse mutant phenotypes by disrupting the functions of discrete genes, likely through the creation of indels and chromosomal rearrangements that result from non-homology directed repair mechanisms such as NHEJ (Sekelsky 2017). However, for many other organisms that were brought into the lab for potential genetic study in the early part of the 20^th^ century, visible mutants were extremely difficult to obtain via mutagenesis by ionizing radiation, suggesting potential differences in the impacts of radiation on genome stability (Fabergé 1983). As such, these organisms did not rise to the level of study seen for the more easily mutable *Drosophila*.

Here we consider the genetic response to ionizing radiation in the dark-winged fungus gnat *Sciara coprophila* (syn. *Bradysia coprophila* and *Bradysia tilicola*). *S. coprophila* has long been of interest among many geneticists due to its unique chromosome biology, which features multiple rounds of programmed chromosome elimination in the germline and in the early embryo, among other events (Gerbi 1986). *S. coprophila* was first cultured as a lab organism by C.W. Metz in the early 1920s (Metz 1925). In the following decades when the mutagenic effects of ionizing radiation were of intense interest, several reports noted that phenotypic changes were exceedingly difficult to induce in *S. coprophila* via ionizing radiation of germline cells, particularly in comparison to *Drosophila* (Smith-Stocking 1936; Metz and Boche 1939; Crouse 1949, 1961; Fabergé 1983). This led to speculation that *S. coprophila* may possess some mechanism of radiation resistance, at least in the germline. More recently, the *S. coprophila* genome was sequenced and its genes annotated (Urban *et al.* 2021), enabling an opportunity to study the radiation response at the level of molecules and gene expression.

To better understand how *S. coprophila* responds to ionizing radiation, we first exposed animals at different developmental stages to varying doses of gamma-irradiation. Our analysis showed the highest resistance for pupae and the least for embryos, consistent with analyses of other organisms. Furthermore, we found that irradiated larvae are able to continue their development after exposures of 80 Gy, which triggers a developmental delay of several days prior to pupation, suggesting an inherent plasticity in the developmental program. Finally, we performed differential gene expression analysis on transcriptomes derived from irradiated and unirradiated larvae, providing a rich dataset for exploration of potential mechanisms of the *S. coprophila* radiation response.

## Materials and Methods

### Fly husbandry

The dark-winged fungus gnat goes by several synonymous names, including *Sciara coprophila* (Lintner 1895)*, Bradysia coprophila* (Steffan 1966), *Sciara/Bradysia tilicola* (Loew 1850), *Sciara amoena* (Winnertz 1867), and others. It is referred to in this manuscript as *S. coprophila* for the sake of continuity with the long historical record of chromosomal, genetic, and molecular research under this name beginning in the 1920s (Metz 1925). All fungus gnats analyzed in this study were female *S. coprophila* strain 7298 Holo2 that descends from the original wavy stocks cultured by Metz and colleagues (Metz and Smith 1931), and that was recently used for genome sequencing and gene annotation (Urban *et al.* 2021). Stocks were obtained from the International Sciara Stock Center at Brown University (https://www.brown.edu/research/facilities/sciara-stock/).

*S. coprophila* adult females predictably produce either all female offspring (gynogenic; wavy wing phenotypic marker) or all male offspring (androgenic; normal wings), permitting simple sexual selection of offspring at all developmental stages. Cultures were maintained at 21°C in a humidified chamber in vials or petri dishes containing a 2.2% agar substrate. Larvae were fed every other day with a dry mix of 2 parts ground shiitake mushroom, 1 part spinach powder, 1 part nettle powder, 4 parts ground oat straw, and 2 parts dry yeast.

### Irradiation and scoring

To collect embryos for irradiation, gynogenic female flies were immobilized on an agar plate and injured around the midsection to induce egg-laying. This method allows for nearly synchronous embryo age upon collection. Embryos were aged 12-14 hours post-egg-laying at the time point of irradiation. To collect 1^st^ instar larvae, gynogenic female and male adults were mass mated to produce vials with a large number of newly hatched female larvae, and agar plugs were extracted from the vials and moved to agar petri plates for irradiation 1-2 days post hatching. All 4^th^ instar larvae and pupae were collected according to their morphology from the progeny of gynogenic females.

Animals were irradiated with ^137^Cs γ-rays from a JL Shepherd irradiator. Doses are given in Gray (Gy), which corresponds to the absorption of 1 J/kg, where 1 Gy = 100 rads. A continuous dose rate of 1.7 Gy/min was administered to animals supported on agar petri dishes enclosed in the chamber. Following irradiation, viability and developmental progression for each stage were scored under a dissecting microscope. Larvae were scored as viable if they showed independent movement during observation. Pupae were scored as viable if they showed movement when prodded with a brush and/or had healthy amber coloration. Pupae that were stiff and dark brown and/or visibly covered with mold were scored as non-viable.

### Library preparation and sequencing

Roughly 590 female 4^th^ instar pre-eyespot larvae (21-28 days post-mating) were divided among three replicate groups. Larvae originating from different mating vials were evenly distributed among the three replicates to reduce potential batch effects from different cultures. Each replicate group was subsequently divided such that half of the larvae were irradiated with 80 Gy over the course of approximately 50 minutes and the other half remained unirradiated. Approximately 45 minutes following radiation treatment, larvae were snap-frozen, and total RNA was extracted and cDNA libraries were prepared as previously described (Urban *et al.* 2021). Libraries were sequenced to yield 100 bp paired-end reads using the Illumina HiSeq 2000 platform.

### Differential Expression and Gene Ontology (GO) enrichment analyses

After inspecting Illumina data using FastQC, quantification of transcripts for each control and irradiated RNA-seq replicate was carried out with Salmon v. 1.2.1 (Patro *et al.* 2017) using an index built on the transcript database generated by the *S. coprophila* genome project (Urban *et al.* 2021) with no decoys. Differential expression between irradiated and non-irradiated larvae was carried out using DESeq2 v. 1.26.0 (Love *et al.* 2014). For each *S. coprophila* gene that exhibited a significant change in gene expression between irradiated and control samples (Benjamini-Hochberg adjusted p < 0.05), the corresponding predicted protein sequence (Urban *et al.* 2021) was compared to the *D. melanogaster* proteome using BLAST (Altschul *et al.* 1990). A *D. melanogaster* protein was considered a potential homolog if it had an e-value less than 1×10^-4^. A reciprocal BLAST was performed with each “hit” *D. melanogaster* protein sequence against the *S. coprophila* proteome; a *D. melanogaster* protein was considered a bona fide homolog if the same *S. coprophila* protein that identified the *Drosophila* protein was subsequently identified as the best reciprocal BLAST hit. Since the *S. coprophila* proteome has not yet been fully functionally characterized, GO enrichment analysis (Ashburner *et al.* 2000) was performed with all *D. melanogaster* hits against the complete *D. melanogaster* gene set using PANTHER v. 16.0 (Mi *et al.* 2019) via http://geneontology.org and the GO-Slim Biological Process output format.

### Phylogenetic analysis

To determine relationships between *S. coprophila* PARP and Argonaute proteins, representative proteins were selected from previous phylogenetic analyses of PARP (Citarelli *et al.* 2010) and Argonaute (Lewis *et al.* 2016) families, and their sequences were obtained from public databases (NCBI Resource Coordinators 2018; UniProt Consortium 2021). PARP and PIWI domain sequences from each protein, including candidate *S. coprophila* homologs, were identified and extracted using Pfam (Finn *et al.* 2014), InterPro (Blum *et al.* 2021), and/or NCBI’s Conserved Domain Database (Lu *et al.* 2020; Blum *et al.* 2021). Phylogenetic analysis was performed on extracted domain sequences using the Phylogeny.fr suite of tools (Dereeper *et al.* 2008), including alignment via MUSCLE, removal of poorly aligned positions and divergent regions via Gblocks, construction of phylogenies using maximum likelihood via PhyML, and rendering of phylogenetic trees via TreeDyn.

## Results

### S. coprophila larvae undergo developmental delay in response to ionizing radiation

To explore how *S. coprophila* survival and development are impacted by ionizing radiation, we exposed animals of various developmental stages to a range of gamma-irradiation dosages. Wildtype *S. coprophila* adults typically live for fewer than five days, and therefore the impact of irradiation on adult survival would be difficult to differentiate from the natural lifespan. Therefore, we restricted our initial analysis to embryonic, larval, and pupal stages.

Embryos aged 12-14 hours after egg-laying were highly sensitive to all doses of gamma-radiation tested (40 - 650 Gy), with no living larvae hatching from irradiated embryos at any irradiated treatment (Table 1). In contrast, irradiated 1^st^ instar larvae survived at rates similar to unirradiated controls up to a dosage of 80 Gy, whereas higher doses resulted in decreased larval viability with reduced size and motility observed for surviving larvae. Similarly, irradiated pupae showed decreased survival at increasing doses of irradiation, although 4/15 (27%) of pupae irradiated at the highest tested dose of 650 Gy were still able to develop and eclose as adults (Table 1).

**Table 1.** Survival of *S. coprophila* at different developmental stages in response to ionizing radiation.

| **Dose (Gy)** | **Embryos irradiated** | **Embryos hatched<sup>1</sup>** | **First instar larvae irradiated** | **First instar larvae survival (14-15d)<sup>1</sup>** | **Pupae irradiated** | **Pupae eclosed** |
| --- | --- | --- | --- | --- | --- | --- |
| 0 | ~200 | ~50-80% | ~50-80 | >50% | 40 | 85% |
| 40 | ~200 | 0 | ~50-80 | >50% | 35 | 80% |
| 80 | ~200 | 0 | ~50-80 | >50% | 35 | 71% |
| 160 | ~200 | 0 | ~50-80 | <50% | 35 | 69% |
| 300 | ~200 | 0 | ~50-80 | <50% | 15 | 73% |
| 650 | ~200 | 0 | ~50-80 | <10% | 15 | 27% |

<sup>1</sup>Estimated proportion of surviving embryos or larvae are based on widefield views of egg-laying surface

We noticed that the larvae that survived high levels of radiation remained atypically small rather than undergoing the rapid growth observed for unirradiated larvae, suggesting a developmental delay or arrest in response to radiation. To further explore this, we exposed larvae of later developmental stages, primarily 4^th^ instar pre-eyespot, to varying doses of irradiation and tracked their developmental progress through pupation and adulthood (Table 2). At 80 Gy, the lowest dose tested, 100% of larvae successfully pupated, whereas the rate of adult eclosion was reduced from 45% in unirradiated controls to 20% in the irradiated set. In contrast, higher levels of radiation led to a decrease in the number of larvae reaching pupation, with exposure from 650 to 1100 Gy resulting in pupation of only 20-30% of larvae even after 3-4 weeks post-treatment. Furthermore, high levels of larval irradiation generally caused complete pupal arrest and/or lethality for those animals that successfully formed pupae, with almost no adult flies emerging from trials using exposure levels greater than 80 Gy (Table 2). Thus, exposure to levels of radiation beyond 80 Gy results in developmental arrest or lethality of *S. coprophila* larvae.

**Table 2.** Arrested development of 4^th^ instar larvae in response to high dosage of ionizing radiation.

| **Dose (Gy)** | **Larvae Irradiated** | **% Pupation<sup>1</sup>** | **% Adult Eclosion<sup>1</sup>** |
| --- | --- | --- | --- |
| 0 | 20 | 100% | 45% |
| 80 | 60 | 100% | 20% |
| 100 | 30 | 97% | 0% |
| 120 | 30 | 87% | 3% |
| 140 | 30 | 60% | 0% |
| 160 | 30 | 23% | 0% |
| 300 | 30 | 27% | 0% |
| 650 | 30 | 17% | 0% |
| 900 | 30 | 27% | 0% |
| 1000 | 30 | 37% | 0% |
| 1100 | 30 | 30% | 0% |

<sup>1</sup>Percentages based on total irradiated larvae for each dose. Larvae were roughly 90% pre-eyespot and 10% eyespot stages.

Although 100% of 4^th^ instar larvae exposed to 80 Gy of gamma-irradiation successfully pupated, the timing of the transition from larva to pupa appeared to be delayed relative to unirradiated controls. To quantify this developmental delay more precisely, we exposed pre-eyespot 4^th^ instar larvae to varying levels of gamma-irradiation and scored the proportion of animals in each of the larval, pupal, and adult stages over time (Figure 2). By Day 6 post-treatment, half of un-irradiated control larvae had transitioned to the pupal stage, while 95% of larvae irradiated with 80 Gy remained in the larval stage. By Day 17, the majority (85%) of animals in the un-irradiated control had emerged as adults, whereas most (85%) animals irradiated with 80 Gy were still in the pupal stage. Consistent with our earlier observations, a higher dose of radiation (160 Gy) resulted in complete developmental arrest in the larval stage for most animals (85%) through day 17, while an intermediate dose of radiation (40 Gy) showed a less severe developmental delay relative to the 80 Gy treatment (Figure 2). In sum, our data support that *S. coprophila* larvae are resistant to doses of ionizing radiation up to 80 Gy, which triggers a developmental delay of approximately 5-8 days prior to pupation.

**Figure 1.**
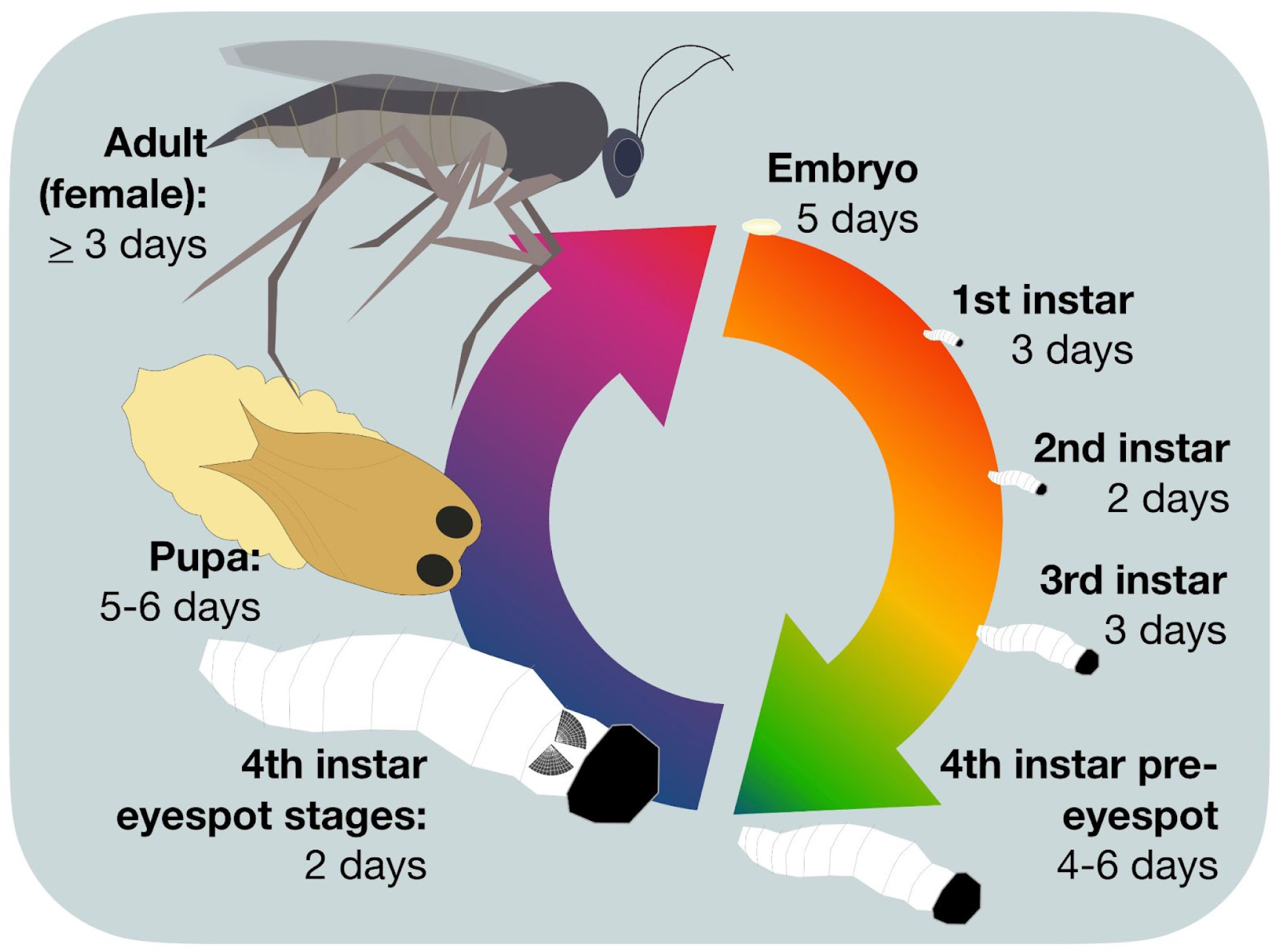
Developmental life cycle of *S. coprophila*. Timing for each stage is for 20°C and corresponds to that reported by Rieffel and Crouse (1966) and our own observations.

**Figure 2.**
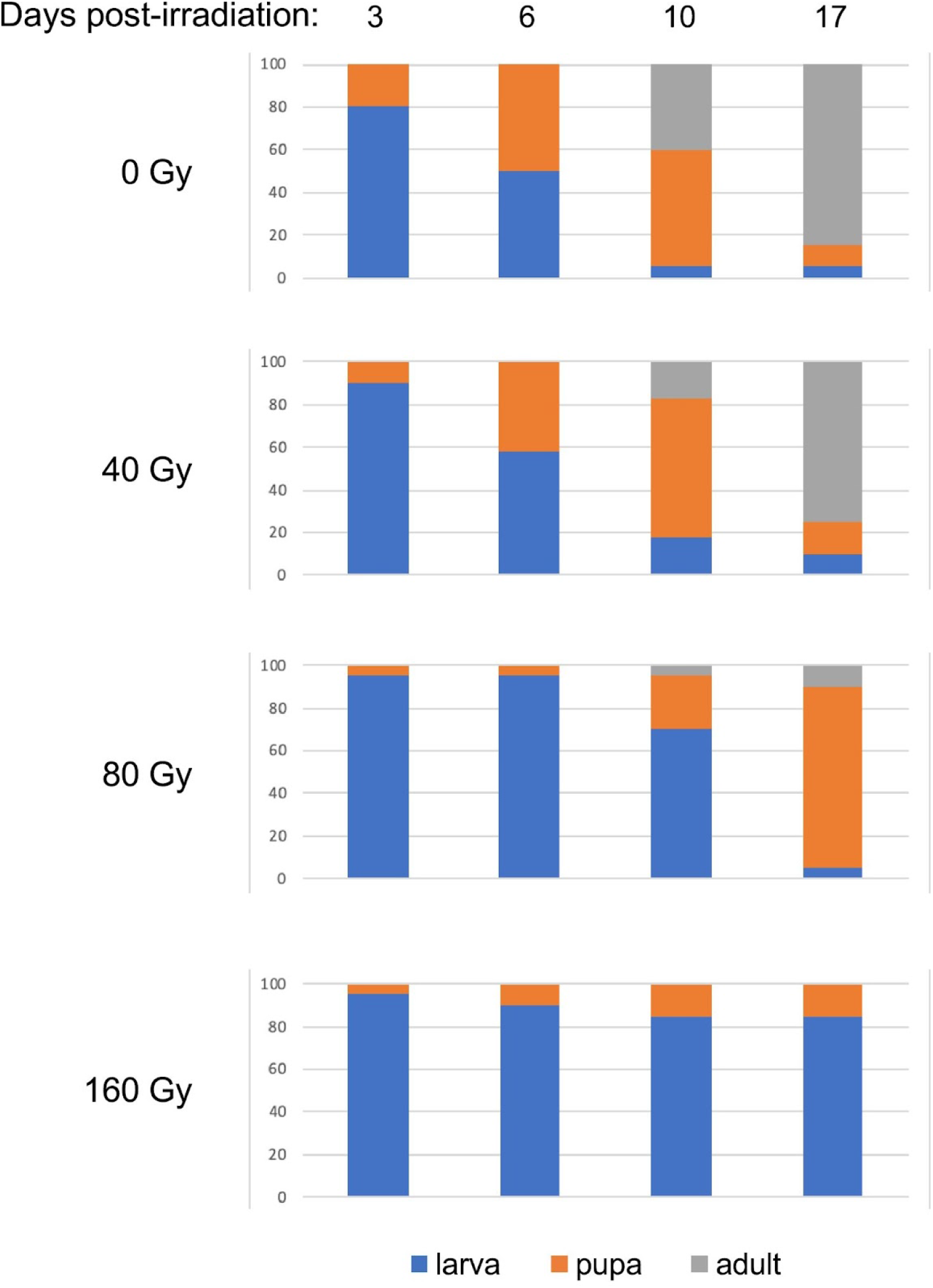
Developmental delay of 4^th^ instar larvae in response to moderate dosage of ionizing radiation. Groups of 20-30 larvae were exposed to the indicated doses of ionizing radiation and developmental progress was followed over the next 17 days. Graphs indicate the percentages of larvae, pupae, and adults for each treatment at each timepoint.

### S. coprophila larvae upregulate DNA repair pathways and downregulate developmental regulators in response to ionizing radiation

To determine the genetic response to ionizing radiation in *S. coprophila*, we divided female pre-eyespot 4^th^ instar larvae into three replicate groups and exposed half of each group to 80 Gy of ionizing radiation, with the remaining half of each group serving as no irradiation controls. We then generated Illumina cDNA libraries to produce between 7 and 10 million paired-end reads for each sample, and subsequently used a bioinformatics pipeline with software packages Salmon (Patro *et al.* 2017) and DESeq2 (Love *et al.* 2014) to perform differential expression analysis between irradiated and non-irradiated groups.

Our analysis showed that 327 of the 23,117 candidate *S. coprophila* genes in the current genome annotation showed significant changes in their expression in response to irradiation, with 232 genes upregulated and 95 genes downregulated (Figure 3A, Supplemental Table 1). To better understand the functions of these genes, we used BLAST searches to identify candidate homologs of each *S. coprophila* protein in the well-characterized *D. melanogaster* proteome. Of the 327 *S. coprophila* genes identified in our experiment, 275 (84.1%) of their protein sequences identified a candidate homolog in *Drosophila* (Supplemental Table 1). We then used those 275 *Drosophila* protein sequences to perform reciprocal BLAST searches against the *S. coprophila* proteome, and found that 195 *Drosophila* proteins returned the same reciprocal best BLAST hit from *S. coprophila*, strongly suggesting an orthologous relationship for each of those proteins (70.9% of hits with homologs; 59.6% of all hits). The remaining 80 proteins that did not return the same reciprocal best BLAST hit are likely members of more complex families with multiple paralogs in at least one of the two species.

**Figure 3.**
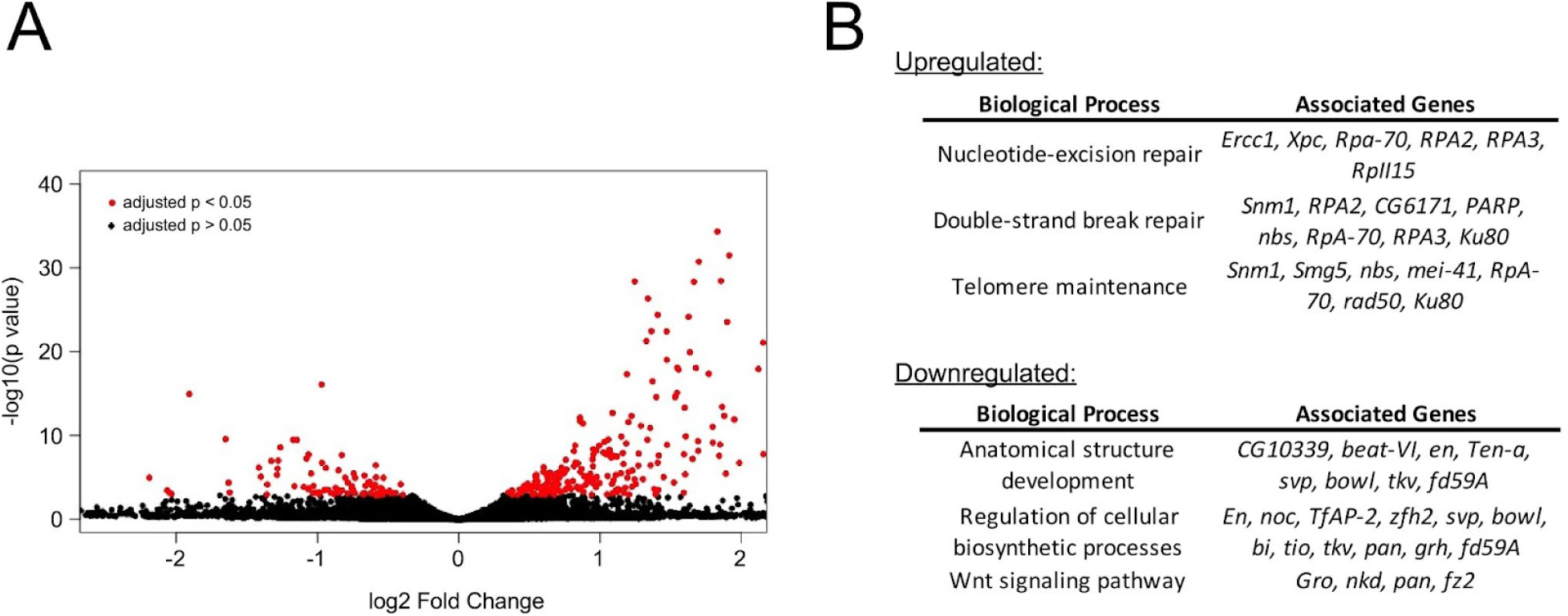
Differential expression analysis of irradiated *S. coprophila* larvae. **A**, volcano plot showing log2-fold change in expression vs −log10 of the adjusted p value for each gene in the study. Points colored red represent genes with significant (p < 0.05) changes in expression. **B**, examples of Gene Ontology (GO) Biological Processes that are enriched among significantly upregulated (top) or downregulated (bottom) genes.

We then used the lists of *Drosophila* homologs identified in our BLAST searches as a proxy for our RNA-seq data to perform GO enrichment analyses for the biological pathways that exhibited changes in expression in response to irradiation in *S. coprophila*. Consistent with the developmental delay observed among larvae irradiated with 80 Gy, we found downregulation of genes involved in Development of Anatomical Structures, Cell Differentiation, Regulation of Cellular Biosynthetic Processes, and Signaling by Wnt Family Regulators (Figure 3B). Furthermore, several genes that are normally upregulated during the late larval and early pupal stages of *Drosophila* development (Graveley *et al.* 2011) were among our *S. coprophila* downregulated gene set, including homologs of *Crp67B* and *ebony*, which are required for development and pigmentation of the *Drosophila* pupa case, respectively (Brehme 1941; Larkin *et al.* 2021), *Samuel* (also known as *Moses*), encoding a co-receptor for nuclear hormone receptor 78 that is upregulated in response to the hormone ecdysone (Baker *et al.* 2007), and *narrow*, which is required for growth of the *Drosophila* wing disc (Ray *et al.* 2015).

In contrast to the downregulated gene set, the genes that were upregulated following irradiation were enriched for GO terms related to DNA repair, including Response to Radiation, Nucleotide-Excision Repair, Double-Strand Break Repair, and Telomere Maintenance (Figure 3B), indicating a robust repair response to radiation-induced DNA damage. Several of the top responding genes were homologs of genes that are important in many human cancers, including poly-ADP-ribose polymerase (PARP) (Slade 2020), Guanine nucleotide binding protein like 1 (GNL1) (Krishnan *et al.* 2020), and Growth Arrest and DNA Damage-inducible 45 (Gadd45) (Tamura *et al.* 2012). Below, we further characterize *S. coprophila* homologs of two notable gene families, PARPs and Argonautes, that are present among the upregulated genes list and represented by large families in the *S. coprophila* genome.

### The S. coprophila radiation response upregulates representatives of PARP and Argonaute gene families

The *S. coprophila* gene *Bcop_v1_g007065* showed by far the largest degree of upregulation (∼68-fold) in response to irradiation. Searches for protein domain signatures in the coding region of *Bcop_v1_g007065* indicated that it has a PARP catalytic domain as well as a series of Ankyrin repeats and a WGR domain. PARP family proteins are known to act in diverse cellular pathways, including DNA damage repair, signal transduction, apoptosis, and chromatin remodeling (Jubin *et al.* 2016). Prior phylogenetic analysis has divided the PARP gene family into six clades with varying catalytic activities and functions, and two of these Clades (Clade 1 and Clade 4) contain members from other species that encode Ankyrin repeats, whereas Ankyrin repeats are not found in known members of the other four clades (Citarelli *et al.* 2010).

To place *Bcop_v1_g007065* among the known PARP family clades, we first carried out a phylogenetic analysis using extracted PARP catalytic domain protein sequences from several representatives of Clade 1 and Clade 4 from different species. Our analysis shows clear placement of *Bcop_v1_g007065* in Clade 1, which includes the canonical human PARP1 that is an important regulator of DNA damage repair, and not in Clade 4, which is comprised entirely of the Ankyrin repeat-rich PARP proteins known as Tankyrases (Figure 4A). Furthermore, two other Ankyrin repeat-rich PARP genes that group with Clade 1, *pme-5* from *Caenorhabditis elegans* and *Adprt3* from *Dictyostelium discoideum*, share the overall domain layout of *Bcop_v1_g007065* with a WGR domain encoded between the Ankyrin and PARP domains, whereas the Clade 4 Tankyrases instead encode a SAM domain in this position (Figure 4B) (Gravel *et al.* 2004; White *et al.* 2009; Citarelli *et al.* 2010; Perina *et al.* 2014). We therefore conclude that Bcop_v1_g007065 is likely a functional ortholog of Pme-5 and Adprt3, and propose the name *<u>P</u>ME-5/<u>A</u>dprt3-<u>L</u>ike <u>P</u>ARP <u>1</u>* (*Bcop-PALP1*) for this highly radiation responsive gene in *S. coprophila*.

**Figure 4.**
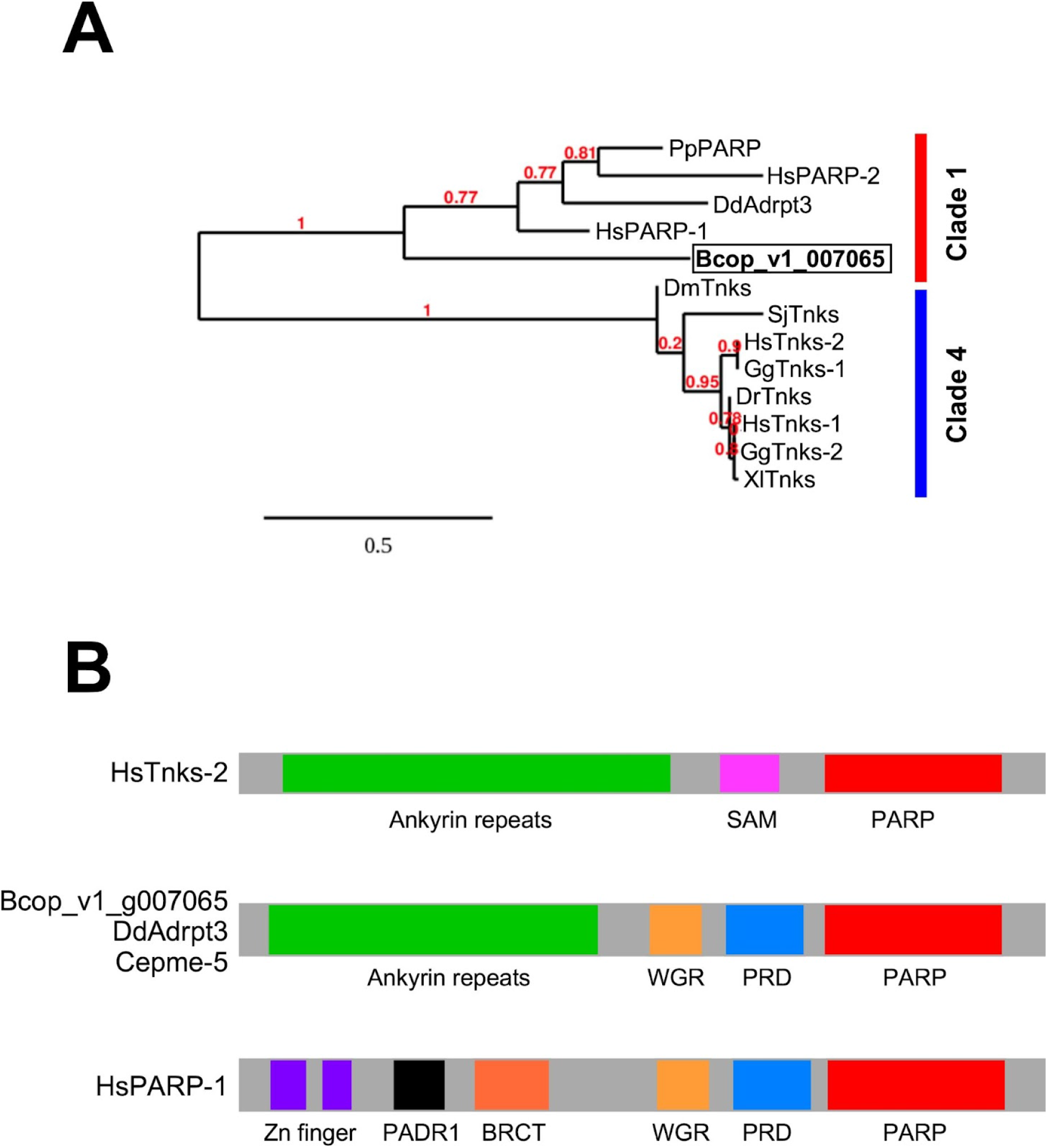
Bcop_v1_007065 (BcopPALP1) is a Clade 1 PARP homolog related to *C. elegans* Pme-5 and *D. discoideum* Adprt3. **A**, phylogenetic analysis of Bcop_v1_007065 PARP domain with representative sequences from Clade 1 and Clade 4 PARPs. Pp, *Physcomitrella patens*, Hs, *Homo sapiens*, Dd, *Dictyostelium discoideum*, Dm, *Drosophila melanogaster*, Dr, *Danio rerio*, Sj, *Schistosome japonicum*, Gg, *Gallus gallus*, Xl, *Xenopus laevis*. **B**, protein domains of *H. sapiens* Tankyrase-2 (Clade 4), Bcop_v1_007065/BcopPALP1, *D discoideum* Adprt3, *C. elegans* pme-5, and *H. sapiens* PARP-1 (Clade 1). SAM, sterile alpha domain, WGR, Trp/Gly/Arg conserved domain, PRD, PARP regulatory domain, PADR1, domain of unknown function conserved in PARP proteins, BRCT, BRCA1 C-terminal domain.

The current genome annotation of *S. coprophila* contains sixteen other candidate genes with homology to the PARP catalytic domain, although only *Bcop-PALP1* showed significant upregulation in response to radiation in our RNA-seq data. To further support our characterization of *Bcop-PALP1*, we carried out a larger scale phylogenetic analysis on all seventeen potential *S. coprophila* PARP homologs using representative sequences of all six Clades from other species (Supplemental Figure 1). We found one other Ankyrin repeat-rich homolog (Bcop_v1_g000291) that groups with the Clade 4 Tankyrases, whereas Bcop-PALP1 once again grouped with Clade 1, further supporting an orthologous relationship for Bcop-PALP1 with Clade1 PARP family members that participate in DNA repair (Citarelli *et al.* 2010).

In addition to the strong upregulation of *Bcop-PALP1* in our RNA-seq data, we noted that three of the significantly upregulated *S. coprophila* genes showed homology to Argonaute family genes. Argonaute proteins are small RNA binding molecules that serve as key effectors in RNAi silencing pathways, including gene silencing by miRNAs, siRNAs, and piRNAs (Wu *et al.* 2020). All known dipteran Argonautes encode PIWI and PAZ domains in their protein sequences, and phylogenetic analysis based on their PIWI domains can divide family members into four Clades: Ago1, Ago2, Ago3, and Piwi/Aubergine (Lewis *et al.* 2016). Notably, members of each Clade carry out different functions, with Ago1 and Ago2 proteins participating in miRNA and/or siRNA silencing, and Ago3, Piwi, and Aubergine participating in biosynthesis and effector steps of piRNA silencing (Meister 2013).

To understand which RNAi pathways may be involved in the radiation response of *S. coprophila*, we carried out a phylogenetic analysis of *S. coprophila* Argonaute homologs, including the three genes upregulated in the radiation response (*Bcop_v1_g003309*, *Bcop_v1_g013021*, and *Bcop_v1_g004567*) along with ten other potential Argonaute homologs identified by the *S. coprophila* genome project (Urban *et al.* 2021) that did not significantly change expression in response to radiation. Using extracted PIWI domains from these proteins and from Dipteran Argonaute homologs that had previously been characterized through phylogenetic analysis (Lewis *et al.* 2016), we found that the three *S. coprophila* Argonautes that were upregulated in response to radiation each grouped with a different Clade, namely Ago1, Ago3, and Piwi/Aubergine, implicating miRNA/siRNA and piRNA pathways in the radiation response (Figure 5). In total, the *S. coprophila* genome encodes multiple representatives of each of the four clades, demonstrating the potential for diverse small RNA pathways in the organisms’ unique biology.

**Figure 5.**
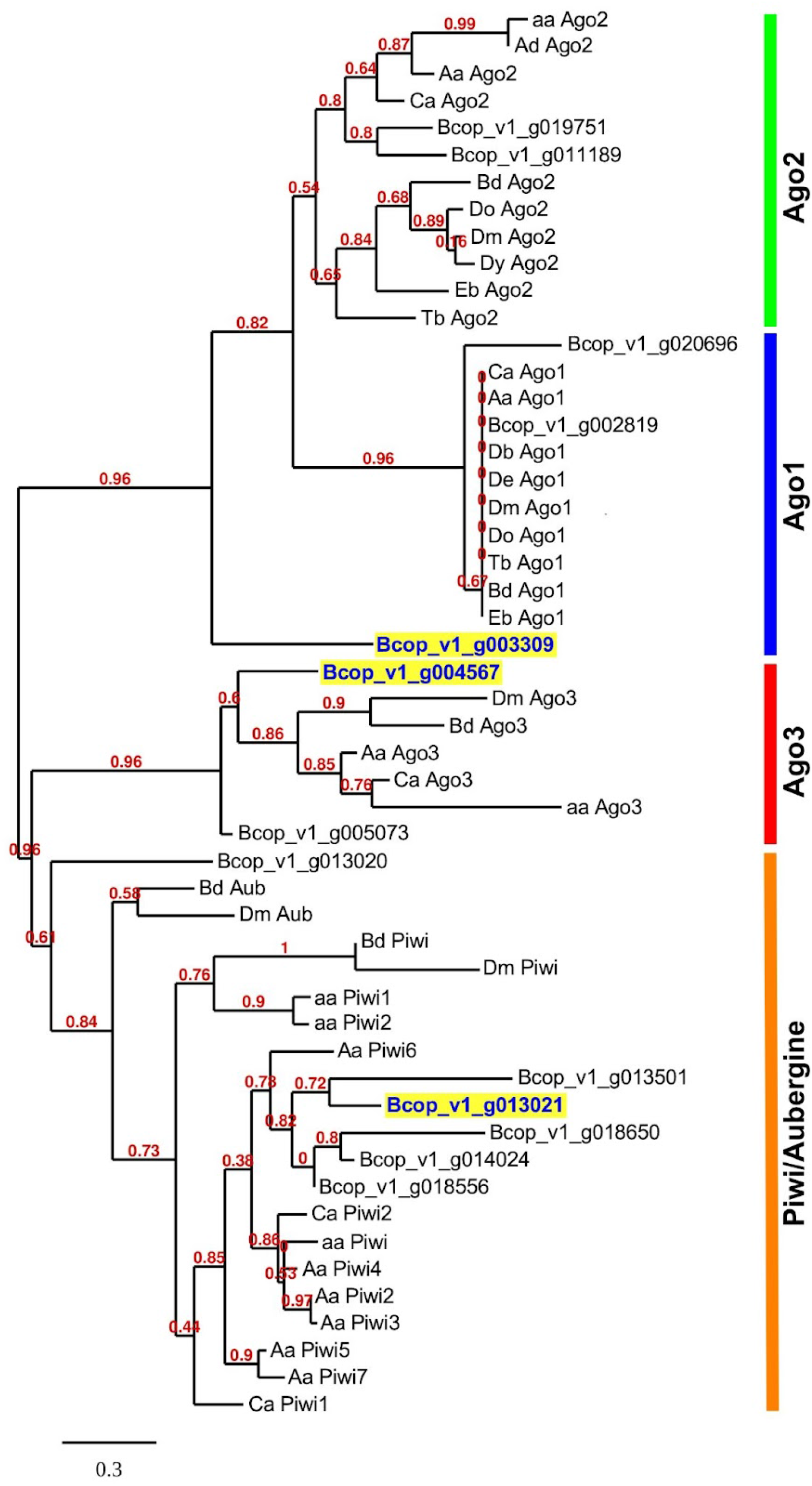
Phylogenetic analysis of Argonaute homologs in *S. coprophila*. Genes encoding Argonaute homologs that are significantly upregulated in irradiated larvae are highlighted. Note that Bcop_v1_003309 groups outside of the Ago1 clade in this tree, but is placed within the Ago1 clade in an expanded analysis of Ago1 and Ago2 sequences (Supplemental Figure 2). Do, *Drosophila obscura*, Tb, *Tabanus bromius*, Db, *Drosophila busckii*, Eb, *Episyrphus balteatus*, Dm, *Drosophila melanogaster*, Ca, *Corethrella appendiculata*, Bd, *Bactrocera dorsalis*, aa, *Anopheles albimanus*, De, *Drosophila erectus*, Dy, *Drosophila yakuba*, Ad, *Anopheles darlingi*, Aa, *Aedes aegypti*.

## Discussion

*Sciara coprophila* was established as a model genetic organism roughly a century ago, and yet very few visible mutants have been recovered since then, which may reflect an inherent resistance to DNA damage induced by ionizing radiation. Historically, studies of potential radiation resistance in *S. coprophila* have focused on transmission of visible mutations or chromosomal rearrangements to the progeny of irradiated organisms (Metz and Boche 1939; Metz and Bozeman 1940; Reynolds 1941; Bozeman and Metz 1949; Crouse 1949, 1961). Here, we further characterize the *S. coprophila* response to ionizing radiation at earlier developmental stages. Our data show that developing *S. coprophila* embryos are highly sensitive to even low doses of gamma-irradiation, whereas larvae can withstand up to 80 Gy and still retain their ability to develop to adulthood. Moreover, differential gene expression analysis of transcriptomes derived from larvae treated with 80 Gy of ionizing radiation relative to unirradiated controls shows upregulation of DNA repair pathways and downregulation of several developmental regulators, providing a key dataset to better understand the radiation response of this unconventional model organism.

Our data demonstrate that irradiated *S. coprophila* larvae delay their transition to pupation by approximately 5-8 days in response to 80 Gy of gamma-irradiation, whereas lower doses have little effect on development, and higher doses result in a “suspended” motile larval state for surviving larvae with no evidence of pupation several weeks after the expected transition would take place. Thus, there appears to be a close coordination between developmental progression and the response to radiation, perhaps analogous to the molecular “checkpoint” mechanisms that prevent progression through the cell cycle in response to DNA damage. Note that *S. coprophila* larvae are also able to suspend their development for up to several months in response to cold temperatures (J. Bliss and S.A. Gerbi, unpublished observation), likely reflecting a state of diapause that serves as the organism’s overwintering strategy. It may be that larval responses to temperature and to irradiation share common genetic mechanisms, a possibility that can be explored in the future via detailed analysis of cold-treated organisms.

Consistent with the delayed developmental progression of irradiated larvae, RNA-seq analysis showed reduced expression of genes known to regulate development in other organisms, including several homeodomain- and zinc finger-family transcription factors and components of the Wnt signaling pathway, marking these genes as candidate regulators of late larval and early pupal development in *S. coprophila*. In holometabolous insects, the transition from larval to pupal stages is largely controlled by ecdysteroid hormones, which increase during late larval stages and impact development via signaling through nuclear hormone receptors such as the Ecdysone Receptor (EcR) (Jindra 2019). It is as yet unclear whether the delayed development in irradiated *S. coprophila* larvae reflects changes upstream or downstream of ecdysteroid biosynthesis and signaling, although we observed a roughly 2-fold downregulation of a *S. coprophila* ortholog of *Samuel*, a gene that plays a role in regulating signaling downstream of EcR in *Drosophila* (Baker *et al.* 2007). Our data may therefore provide insight into the molecular connections between *S. coprophila* hormone signaling and developmental effectors.

As expected, ionizing radiation led to the upregulation of genes involved in several DNA repair pathways, including NER and NHEJ, representing a common response to DNA damage among metazoans (Sekelsky 2017). The highest change in expression was observed for a PARP family homolog that we named *Bcop_PALP1*, which is orthologous to *C. elegans pme-5* and *D. discoideum Adprt3* according to overall structure and catalytic domain sequence homology. Although functional characterization of *Adprt3* is lacking, *pme-5* is among the most transcriptionally upregulated genes in response to ionizing radiation in worms, and DNA damage-induced germ cell apoptosis is increased in worms where *pme-5* expression is knocked down via RNAi, providing functional evidence of a role in the DNA damage response (Gravel *et al.* 2004). *Bcop-PALP1* is one of seventeen PARP genes annotated by the *S. coprophila* genome project (Urban et al. 2021), representing a large expansion of this gene family, particularly in comparison to the *Drosophila* genome, which encodes only a single Clade 1 PARP1 homolog and a single Clade 4 tankyrase (Larkin *et al.* 2021). At least six of the *S. coprophila* PARP homologs group within Clade 1, where many members from other species have been shown to have roles in DNA repair (Citarelli *et al.* 2010). The expansion of the PARP gene family may therefore be related to the potential radiation resistance of *S. coprophila*, with *Bcop-PALP1* likely playing a central role.

We also noted three Argonaute homologs among the genes that were significantly upregulated in response to radiation. Previous studies from both plants and animals have shown that double-strand breaks in DNA generate small RNAs that can be bound by Argonaute homologs, which facilitates recruitment of repair factors such as Rad51 (d’Adda di Fagagna 2014; Gao *et al.* 2014; Oliver *et al.* 2014; Rzeszutek and Betlej 2020; Hu *et al.* 2021). *S. coprophila* Argonaute homologs may therefore directly participate in DNA repair pathways. Furthermore, DNA damage is known to activate and mobilize transposable elements (TEs) in both prokaryotes and eukaryotes (McClintock 1984; Bradshaw and McEntee 1989; Walbot 1992; Eichenbaum and Livneh 1998; Rudin and Thompson 2001; Hagan *et al.* 2003; Farkash and Luning Prak 2006). Phylogenetic analysis showed that two of the upregulated *S. coprophila* Argonaute genes are orthologous to Ago3 and Piwi proteins that are key components of the piRNA pathway, which plays an important role in silencing TEs in the germline of metazoan organisms (Wu *et al.* 2020). The upregulation of *S. coprophila* Ago3 and Piwi orthologs may therefore be related to maintaining genome integrity in the germline by preventing mobilization of TEs. It is also possible that upregulation of Argonaute homologs aids in the post-transcriptional regulation of other *S. coprophila* genes through activation of the RNA-Induced Silencing Complex (Pratt and MacRae 2009), perhaps suppressing the translation of other developmental regulators until DNA damage has been repaired. A detailed analysis of small RNAs in irradiated larvae may reveal an additional layer of gene regulation that is not captured by RNA-seq methods.

Finally, our analysis provides a modern update to a rich history of research into potential radiation resistance in *S. coprophila*, largely motivated by the recalcitrance of the organism to the appearance of visible phenotypes in response to radiation. Early studies of *S. coprophila* showed that chromosomal rearrangements are readily isolated from irradiated males, demonstrating that sperm are sensitive to radiation, but rearrangements could not be isolated from irradiated oocytes until the later stages of meiosis beyond Metaphase I, supporting that mechanisms of radiation resistance exist in developing oocytes (Metz and Boche 1939; Metz and Bozeman 1940; Reynolds 1941; Bozeman and Metz 1949; Crouse 1950). However, visible phenotypes are equally difficult to isolate from either irradiated sex (Smith-Stocking 1936), which brings into question whether there is indeed a relationship between radiation resistance and a historical lack of discernable markers. Thus, alternative explanations have been suggested to account for the difficulty in isolating visible phenotypes, including the unique sex-determination system of *S. coprophila* that prevents novel recessive autosomal mutations from being isolated as homozygotes until four generations after irradiation, adding complication to genetic screens (Crouse 1949). Our analysis shows that *S. coprophila* larvae exposed to 40 Gy of gamma radiation show no discernible difference from unirradiated larvae in their developmental progression, whereas exposure to 80 Gy and above does impact development. Although direct comparisons can be difficult to make due to differences in biology, *Drosophila* larvae exposed to 40-50 Gy under very similar conditions to our study show significant reduction in their ability to continue development (Sudmeier *et al.* 2015). *S. coprophila* therefore appears to show heightened resistance to radiation relative to *Drosophila* in this type of assay, although the difference is relatively modest. Ultimately, it may be that *S. coprophila* encodes a set of robust DNA repair pathways that, when combined with other unique aspects of its biology, result in the apparent radiation resistance of the organism.

## Acknowledgements

Thanks to Richard Shea at Brown University for assistance with radiation treatments and to Sebastien Santini (CNRS/AMU IGS UMR7256) and the PACA Bioinfo platform (supported by IBISA) for the availability and management of the phylogeny.fr web tools. JMU was supported by predoctoral traineeships from the NIH (T32 GM007601), NSF/EPSCoR (#1004057) and an NSF predoctoral fellowship (GRFP-DGE-1058262); JRB, KRG, AB, and BJT were supported by grants from the NIH (P20 GM0103423 and R15 GM132896-01); SAG was supported by the NIH (R01 GM121455).

---

**Supplemental Figure 1.**
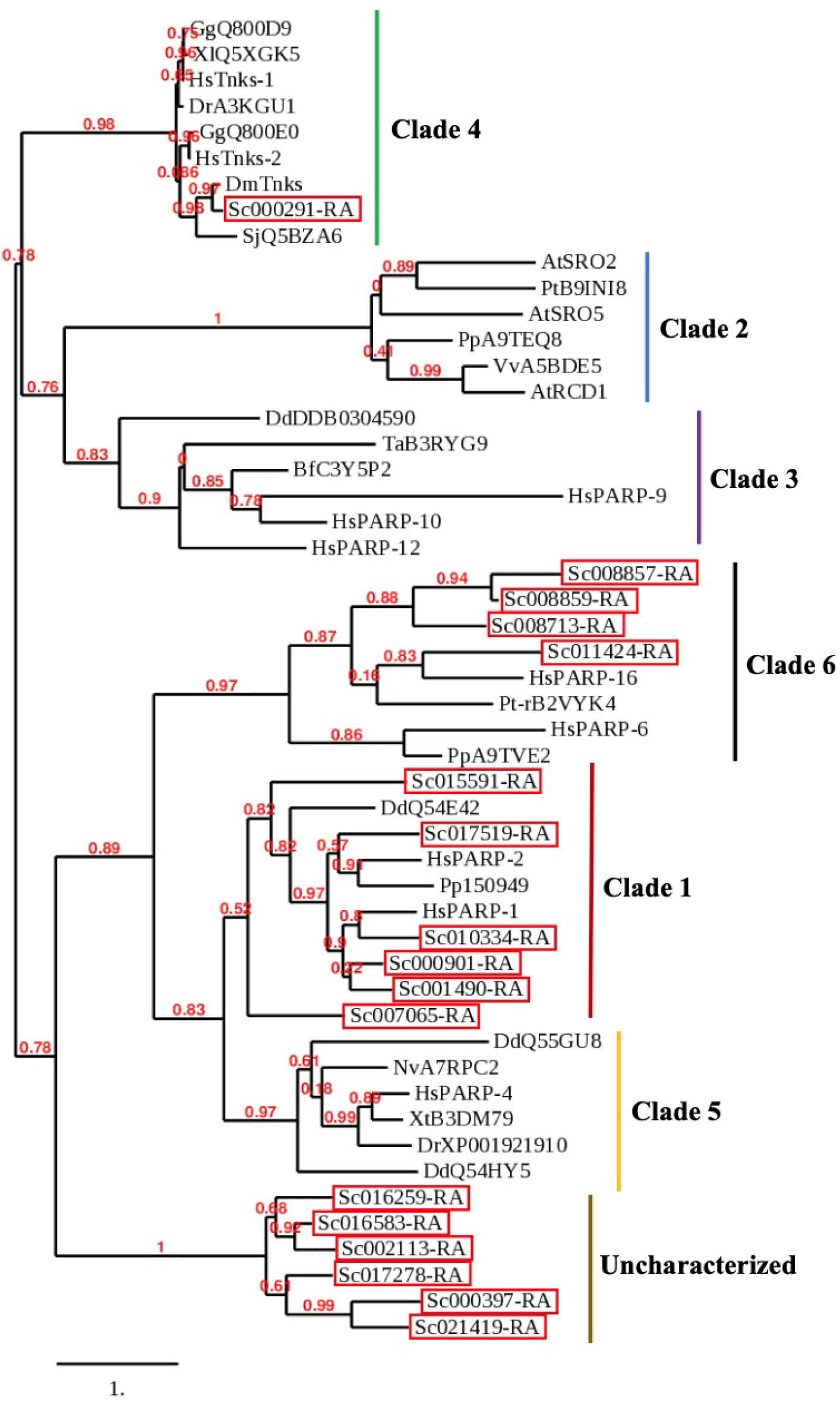
Phylogenetic analysis of PARP homologs in *S. coprophila*. Seventeen annotated *S. coprophila* genes with PARP catalytic domains (Urban *et al.* 2021) were analyzed along with homologs from other species representing all six clades (Citarelli *et al.* 2010). Catalytic domain sequences for representative PARPs were taken from *Physcomitrella patens* (Pp), *Vitis vinifera* (Vv), *Arapidopsis thaliana* (At), *Populus trichocarpa* (Pt), *Homo sapiens* (Hs), *Trichoplax adhaerens* (Ta), *Branchiostoma floridae* (Bf), *Dictyostelium discoideum* (Dd), *Drosophila melanogaster* (Dm), *Schistosome japonicum* (Sj), *Gallus gallus* (Gg), *Danio rerio* (Dr), *Xenopus laevis* (Xl), *Xenopus tropicalis* (Xt), *Nematostella vectensis* (Nv), and *Pyrenophora tritici-repentis* (Pt-r). *S. coprophila* gene names were abbreviated to ScXXXXXX for simplicity (red boxes). All known PARPs were correctly sorted into their published Clades (Citarelli *et al.* 2010). Six *S. coprophila* PARPs group with Clade 1, the DNA repair clade, including Sc007065/Bcop-PALP1; one groups with the Clade 4 tankyrases; four group with the Clade 6, which are likely ancient mono-ADP-ribosyltransferases with roles in membrane biology (Vyas *et al.* 2013). The six remaining S. coprophila PARP homologs were not placed in known clades.

**Supplemental Figure 2.**
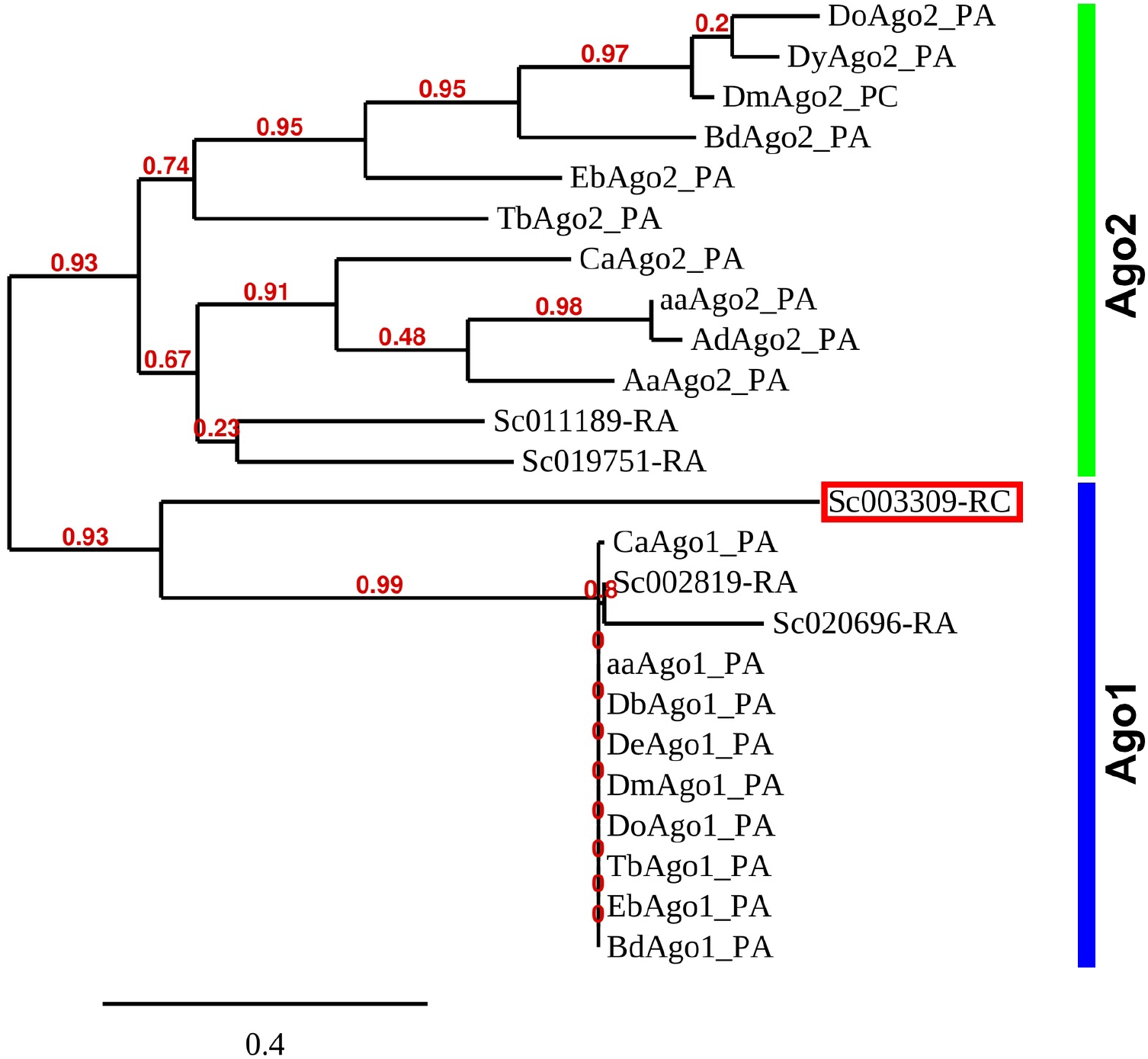
An *S. coprophila* Ago1 ortholog is upregulated in response to radiation. *S. coprophila* Argonaute homologs that grouped with Ago1 or Ago2 clades in Figure 5 were re-analyzed with an expanded pool of Ago1 and Ago2 orthologs from other Dipteran species. Bcop_v1_003309 (simplified here to Sc003309, red box) more confidently groups with the Ago1 clade according to this analysis.

